# Modulation of neural gene networks by estradiol in old rhesus macaque females

**DOI:** 10.1101/2023.12.18.572105

**Authors:** Rita Cervera-Juanes, Kip D. Zimmerman, Larry Wilhelm, Dongqin Zhu, Jessica Bodie, Steven G. Kohama, Henryk F. Urbanski

## Abstract

The postmenopausal decrease in circulating estradiol (E2) levels has been shown to contribute to several adverse physiological and psychiatric effects. To elucidate the molecular effects of E2 on the brain, we examined differential gene expression and DNA methylation (DNAm) patterns in the nonhuman primate brain following ovariectomy (Ov) and subsequent E2 treatment. We identified several dysregulated molecular networks, including MAPK signaling and dopaminergic synapse response, that are associated with ovariectomy and shared across two different brain areas, the occipital cortex (OC) and prefrontal cortex (PFC). The finding that hypomethylation (*p*=1.6×10^-51^) and upregulation (*p*=3.8×10^-3^) of *UBE2M* across both brain regions, provide strong evidence for molecular differences in the brain induced by E2 depletion. Additionally, differential expression (*p*=1.9×10^-4^; interaction *p*=3.5×10^-2^) of *LTBR* in the PFC, provides further support for the role E2 plays in the brain, by demonstrating that the regulation of some genes that are altered by ovariectomy may also be modulated by Ov followed by hormone replacement therapy (HRT). These results present real opportunities to understand the specific biological mechanisms that are altered with depleted E2. Given E2’s potential role in cognitive decline and neuroinflammation, our findings could lead to the discovery of novel therapeutics to slow cognitive decline. Together, this work represents a major step towards understanding molecular changes in the brain that are caused by ovariectomy and how E2 treatment may revert or protect against the negative neuro-related consequences caused by a depletion in estrogen as women approach menopause.

## 1 | INTRODUCTION

Estradiol (E2) is mainly produced by the ovaries and is the most physiologically relevant estrogen (others include estrone and estriol). E2 exerts control over numerous biological functions by binding to the intracellular estrogen receptors alpha and beta (ERα and ERβ) [1–5] and the G-protein coupled receptor GPR30/GPER1 [6].

E2 levels fluctuate throughout the female menstrual cycle and, as women age and go through natural menopause, E2 levels progressively decline. As a reflection of the broad roles of E2, adverse physiological and psychiatric effects accompany the natural decline of E2 levels. These include vasomotor symptoms (hot flashes, night sweats) [7, 8], sleep disturbances [9], somatic symptoms (pains, aches) [7], cognitive performance decline [10], anxiety [7], and depression [7]. Such effects are expected given the E2’s neural role in mediating synaptic plasticity [11–16], increasing dendritic spine density, long-term potentiation (LTP) [17], neuroprotective effects [18] and improving cognitive performance [19, 20].

A lack of E2 right after menopause negatively affects learning and memory and increases the risk of neurodegenerative diseases, such as Alzheimer’s disease (AD) [21, 22]. The incidence of AD and related dementias is two to three times higher in women than in men, and premature menopause increases this risk [23, 24]. Although surrounded by a lot of debate [25, 26], there is much evidence to suggest that E2 replacement therapy, administered immediately at menopause [27–35], may improve cognitive performance and reduce risk for onset and development of AD [3–5, 24]. These effects highlight E2’s role in preserving cognitive function and overall well-being.

Although it does not exactly replicate the natural decline in E2 as seen in healthy women, ovariectomy (Ov) and ovohysterectomy (OvH) have been widely used in preclinical models to investigate the physiological and neural adaptations that take place when the ovarian E2 supply is removed [18]. It should be noted that, although there is neuronal production of E2 with important neuromodulator and neuroprotective functions [36, 37], its source are androgens, which are mostly produced by the ovaries [38]. Ov and Ov-HRT are implemented as effective cancer treatment and are commonly used for benign gynecologic conditions in women 40 years and older [39]. Furthermore, women who underwent Ov showed a higher risk for development of dementia, but not if they received Ov-HRT treatment at the time of surgery [33]. Thus, preclinical animal surgical menopause animal models become excellent models in which to examine the negative impact of reduced E2 concentrations on molecular and physiological processes, as well as the potential benefits of hormonal replacement therapy (HRT), in women who have experienced abrupt E2 removal.

Among the current preclinical models, nonhuman primates (NHPs) are highly valuable for this research because of their very similar physiology to humans and because females undergo a typical menopausal transition [40]. Our own studies, and others, demonstrated that immediate Ov-HRT in aged surgically menopausal rhesus macaque females showed positive effects on memory [30, 41] and favorable effects on cognition in aged females under an obesogenic diet [42]. Using brain samples from the same females, we identified differential gene expression in the occipital (OC), prefrontal cortex (PFC), hippocampus (HIP) and amygdala (AMG), with an enrichment in neuroinflammation in OC and HIP, but an inhibition in the AMG with Ov-HRT. Synaptogenesis, circadian rhythm, mitochondrial dysfunction, mTOR, glutamate, serotonin, GABA, dopamine, epinephrine/norepinephrine, glucocorticoid receptor signaling, neuronal NOS, and amyloid processing were exclusively enriched in AMG. As compared to the control group, most of these signaling pathways are downregulated after Ov-HRT, suggesting a protective effect of E2 in Ov-HRT females under a Western-style diet. A follow up study, using the contralateral AMG from these same females, as well as from a separate cohort of females under a regular chow diet, showed that Ov-HRT (immediate treatment) had lower histological amyloid b plaque density as compared to placebo females [43]. Furthermore, our own studies showed that E2 treatment clearly improved cognitive performance in the same animals included in the current study [30]. In the present study, we sought to elucidate the molecular pathways in two cognitive-relevant cortical regions that could be altering brain function and ultimaltely contributing to such cognitive benefits.

It is well known that E2 binds to ERα and ERβ, and through the canonical mechanism of action, the E2-ER complex binds to estrogen-responsive elements (ERE) at promoters of target genes regulating their expression. In addition, E2, through binding to EREs, regulates gene expression through neuroepigenetic regulation [44]. After binding to ERE, the ligand-bound ERs recruit chromatin remodelers, such are BAF60 or recruiting CREB binding protein (CBP), that regulate DNA and histone modifications [45–49]. Intrahippocampal E2 increases DNMT3a and 3b levels and activity, decreased HDAC2 expression, increased H3 and H4 acetylation, altering memory in ovariectomized mice. Furthermore, DNMTs inhibiton by 5-AZA inhibited recognition memory [50, 51]. These prior findings support a critical role of DNAm in mediating the effects of E2 in brain function.

Iin the present study, we characterize the transcriptomic and methylomic profile of the brain between elderly ovary intact (OI), Ov females without HRT under a regular chow diet to determine the molecular signatures associated with an abrupt depletion of E2 at a peri-menopausal age. We next evaluate if HRT can revert any of these changes to maintain an age-matched molecular profile. We focus on two cortical brain regions associated with cognitive function and known to be impacted in aging and dementias. The OC is involved in visuospatial processing, distance and depth perception, color determination, object and face recognition, and memory formation (Rehman & Al Khalili, 2021; Stufflebeam & Rosen, 2007). Damage in this area is linked to hallucinations in dementia patients. The PFC is a central brain structure involved in working memory, temporal processing, decision making, flexibility, and goal-oriented behavior [52]. In the context of AD, neurodegeneration and neural damage spreads from the PFC to occipital lobes.

The present study focused on elucidating the molecular effects of E2 on the primate brain by examining the differential gene expression and DNAm patterns in OI and following Ov and subsequent E2 treatment. With this model, we have identified a number of dysregulated molecular networks that are associated with Ov and are shared across two different regions of the brain, OC and PFC. The latter being particularly susceptible to age-associated neuropathologies such as AD and frontotemporal lobar degeneration (FTLD). Our results suggest extensive molecular differences in the brain induced by E2 depletion. We have also identified a number of Ov-related molecular differences that appear to be modulated by Ov-HRT treatment. These changes offer valuable insights into the neurobiological consequences of E2 deficiency, and potential alternative therapeutics that could be more targeted.

## 2 | RESULTS

### 2.1 | Differential Expression Analysis

In the OC, 14,842 genes met the filtering criteria while 14,590 genes met the filtering criteria in the PFC. After computing association testing with each gene from each of those sets, we identified 150 and 128 differentially expressed (DE) genes associated with Ov (FDR < 0.05) in the OC and PFC, respectively (Table S1).

To explore if Ov-HRT treatment modulates the effect of Ov, we computed an interaction test between Ov-HRT treatment and Ov status. Instead of globally testing for an interaction between Ov-HRT and Ov status, we were primarily interested in genes that showed significant expression changes related to Ov that no longer showed a statistical association with Ov once the animals received Ov-HRT treatment. To obtain a general set of genes that fit this criterion, we filtered the results to include only genes that showed suggestive evidence of Ov association (unadjusted *p* < 0.05) without Ov-HRT treatment and little evidence of Ov association with Ov-HRT treatment (unadjusted *p* > 0.1). From this reduced set of 884 (PFC) and 663 genes (OC), we further filtered down to only those results that had suggestive evidence (*p* < 0.05) of interaction between Ov and Ov-HRT treatment. In the OC and PFC, we identified 19 and 10 genes, respectively, with suggestive evidence for Ov-HRT effects (Table 1). Ten of these genes are known to interact with the estrogen receptors or its expression being associated with the levels of estrogen. These include the transient receptor potential vanilloid 6 (*TRVP6*), adrenomedullin (*ADM*), the glucose transporter 12 (*SLC2A12*), supervillin (*SVIL*), acyl-CoA synthetase 2 (*ACSF2*), lymphotoxin B receptor (*LTBR*), the hematopoietic PBX-interacting protein 1 (*PBXIP1*), the fucosyltransferase 1 (*FUT1*), neuromedin U (*NMU*) and the Purkinje cell protein 4 (*PCP4*). Furthermore, *TRVP6* is known to contain estrogen-responsive element in its promoter. In the OC-specific network, *ADM* was part of the insulin secretion pathway. And in the OC/PFC combined network, *PBXIP1* was a member of the AMPK signaling pathway.

**TABLE 1.**
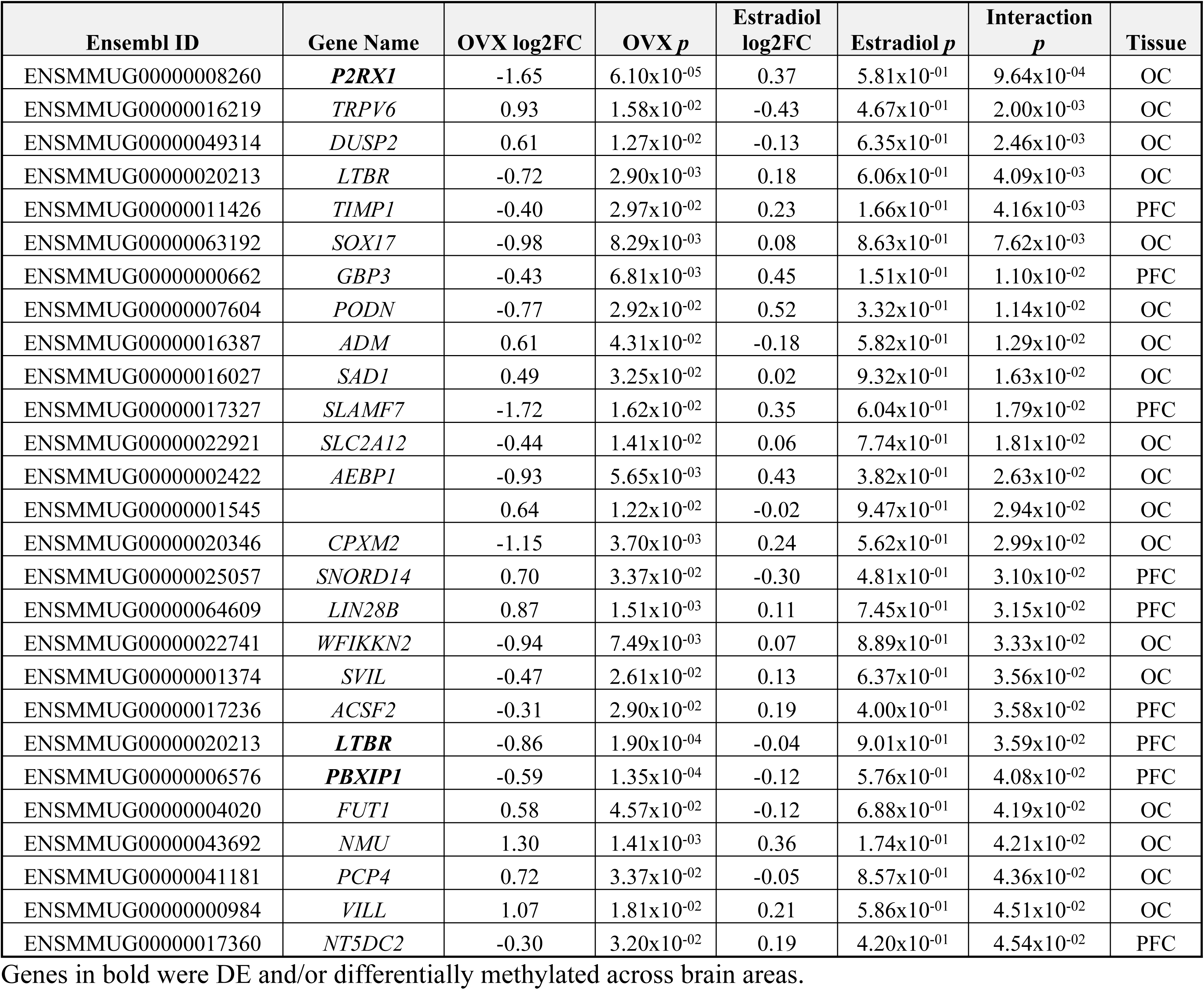
Interaction Results. Genes that demonstrate suggestive evidence (*p* < 0.05) for a modifying E2 result.

### 2.2 | Differential Exon Usage Analysis

These represent a unique set of genes that are being alternatively spliced in Ov animals and are mutually exclusive from the set of DE genes. Among these genes, *HADHB, MDH2* and *ELMO1* are known to interact with ERs and/or have EREs (i.e. *ELMO1*). No exons were identified as significant in the test of interaction between Ov and Ov-HTR. Given the massive number of tests computed, the smaller sample size of the study, and the additional degrees of freedom needed to test the interaction between exon and Ov status (see Methods), it is likely that we are underpowered to compute DEU analysis at this scale. Nonetheless, the 15 unique genes demonstrating significant DEU were included in the network analysis (Figure 1). *BMS1*, *CACNA2D3*, *DAPK1* and *RGS6* clustered into the MAPK signaling and ribosome biogenesis networks (Figure 1). In addition, we overlapped each of the 15 DEU genes with significant DMRs in the OC (FDR<0.05 and Sidak<0.05), because exon usage can often be influenced by DNAm. We did not identify any overlapping results between the genes that were mapped to our significant DMRs and the DEU genes.

**FIGURE 1.**
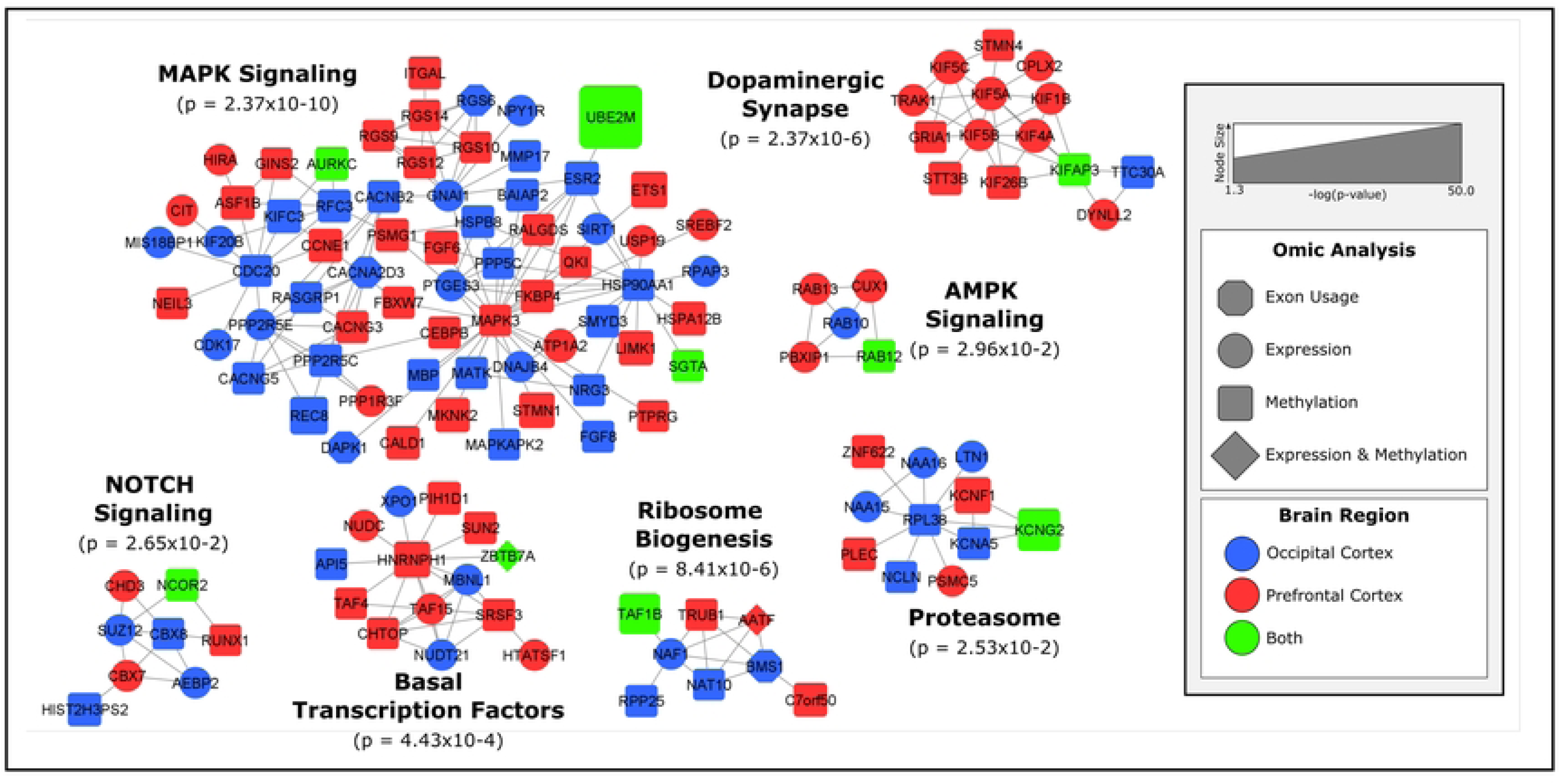
Biological networks of ovariectomy-related changes in gene expression and DNA methylation. Protein interactions were obtained from STRING’s protein interaction database. MCODE was used to find tightly connected clusters of interactions that are labeled according to function defined in Gene Ontology biological processes (enrichment p-value listed). The color of the nodes reflects the tissue where the gene was identified while the shape of the nodes reflect which omics analysis the gene was identified in. The size of the node reflects statistical significance with larger nodes, like *UBE2M*, being more significant.

### 2.3 | Differential Methylation Analysis

In the OC and PFC, 2.6 million and 2.9 million CpGs met the filtering (no missing values across samples and standard deviation of CpG methylation rate across all samples less than 5%), respectively. After computing association testing with each CpG in from each of those sets, and aggregating the CpG results into DMRs, we identified 254 (OC) and 457 (PFC) significant (Sidak’s *p* < 0.05) DMRs associated with Ov (Table S1). 18 DMRs were shared across both brain regions (Table 2). All of them showed the same direction of change in DNAm, except for the E3 ubiquitin protein ligase, *TRIM36,* and the SH3 binding kinase 1, *SBK1*, that were hypomethylated in OC. Furthermore, *SBK1* is both hypermethylated and downregulated in the PFC. In the PFC, 5 DMRs showed overlap with DE genes (Table 2). The *UBEM2* (ubiquitin conjugating enzyme 2) DMR is very strongly replicated across both brain regions, which again suggests this is unlikely to be a false positive (Figure 2). *UBE2M* also shows suggestive evidence of differential expression in the OC (Figure 2). In this case, the DMR (OC: 157 DMCs, PFC: 145 DMCs) is hypomethylated in both brain areas with Ov as compared to OI, and the gene is upregulated in the OC of Ov with or without HRT. The DMR is located ∼3 kb upstream of the *UBE2M* transcription start site, and mapping to the last exon of the *MZF1* gene. Differential expression was detected for *UBE2M* but not for the *MZF1* gene. According to the reporter assay results, the subregion within this DMR has promoter activity (Figure S1), as shown by an increased luciferase relative light units (RLUs) as compared to the control vectors (PGL3 enhancer vector, Figure S1).

**FIGURE 2.**
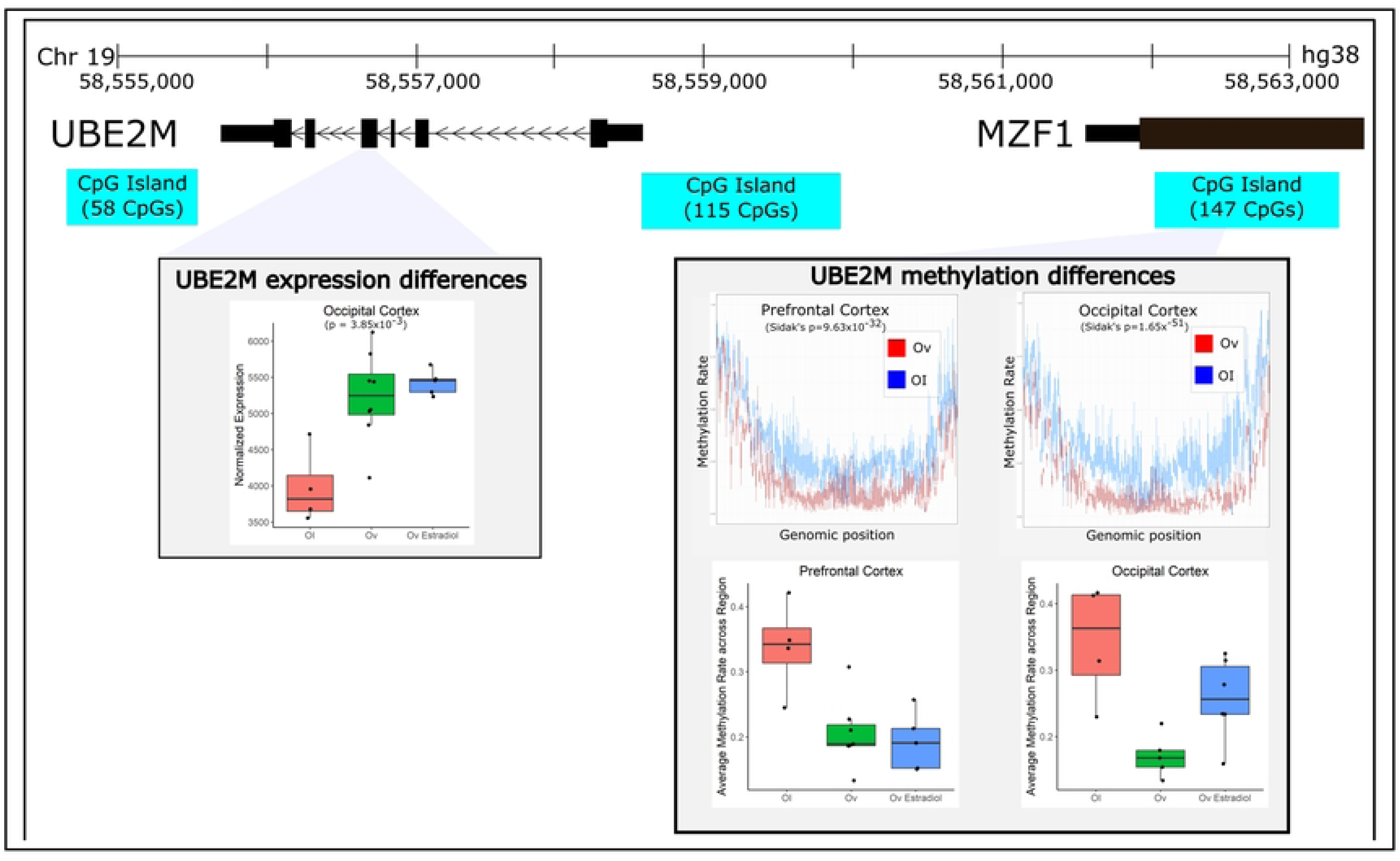
Ovariectomy-related changes in gene expression and methylation of *UBE2M*. *UBE2M* was a gene that demonstrated significant differences in methylation across both brain regions. In addition, it demonstrated significant differences in expression in the OC. The DMR and CpG island that was annotated to *UBE2M* resides upstream of *UBE2M* in the same genomic region as the *MZF1* gene and appears to be hypo-methylated with ovariectomy. Expression of *UBE2M* increases with ovariectomy, regardless of E2 treatment and average DNA methylation rates are not significantly altered with E2 treatment.

**TABLE 2.**
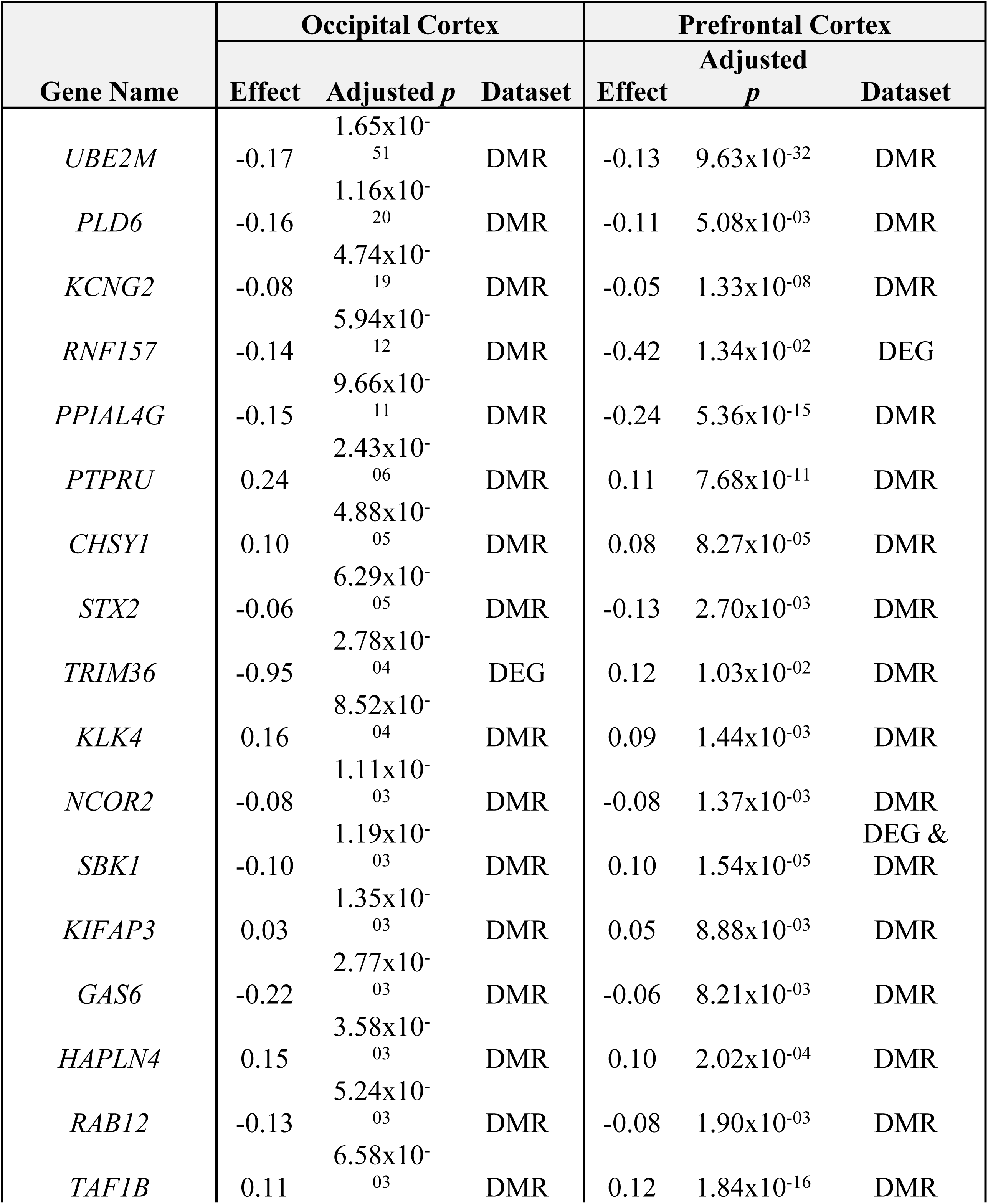
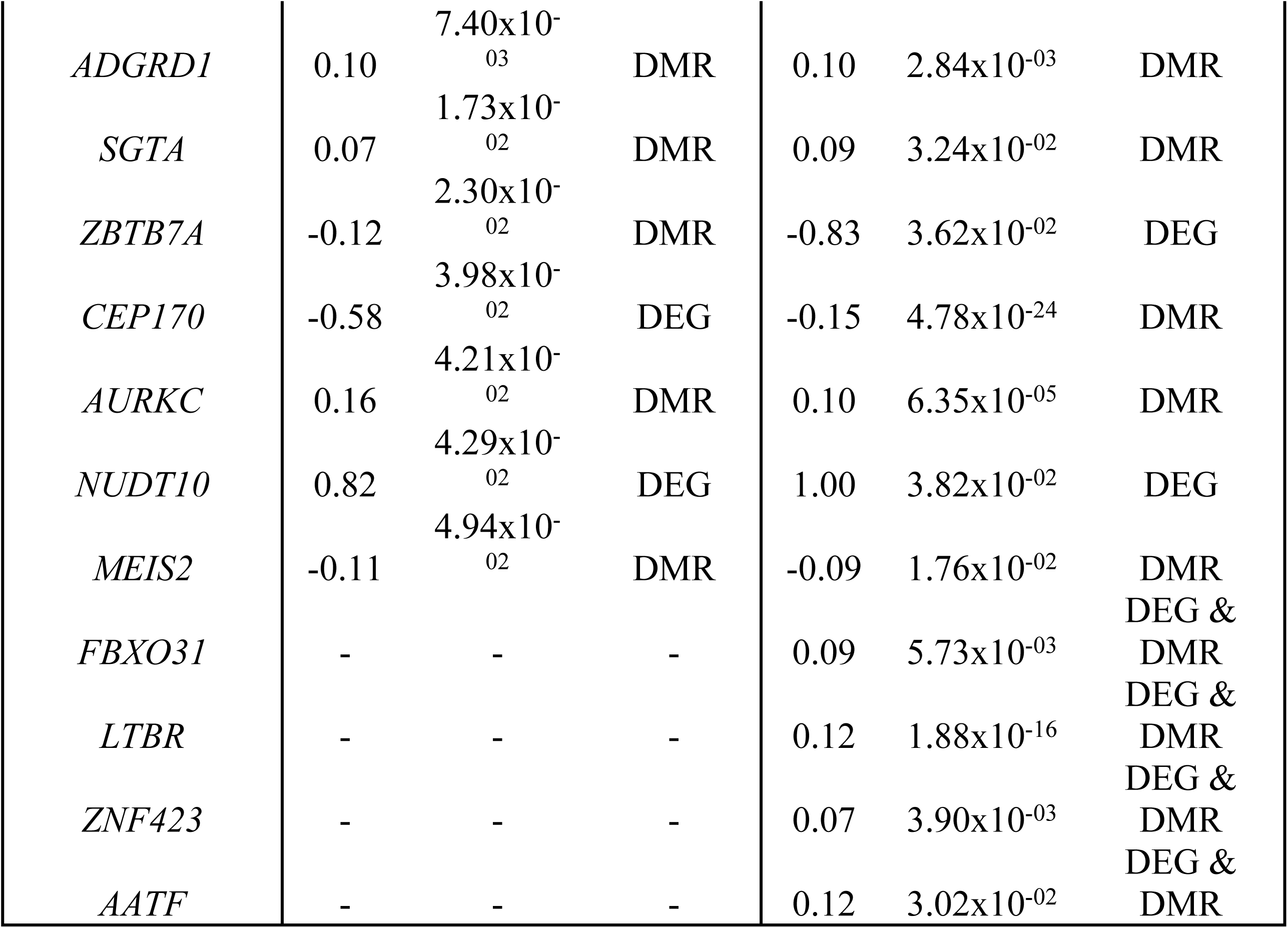
Overlapping Results. Genes that demonstrate either differential expression or methylation across both the occipital cortex and the prefrontal cortex. DMR stands for differentially methylated region and DEG stands for differentially expressed gene. “Effect” stands for either a log_2_(fold-change) or a difference in methylation rates. A positive effect is indicative of an increase with ovariectomy, while a negative effect indicates a decrease with ovariectomy. “Adjusted *p*” is the *p*-value adjusted for multiple comparisons, either an FDR adjustment for DEGs or Sidak’s adjustment for DMRs. For results that are both a DEG & DMR in the same tissue, the DMR effect and p-value are listed.

| Gene Name | Occipital Cortex |  |  | Prefrontal Cortex |  |  |
| --- | --- | --- | --- | --- | --- | --- |
| | Effect | Adjusted $p$ | Dataset | Effect | Adjusted $p$ | Dataset |
| *UBE2M* | -0.17 | $1.65 \times 10^{-51}$ | DMR | -0.13 | $9.63 \times 10^{-32}$ | DMR |
| *PLD6* | -0.16 | $1.16 \times 10^{-20}$ | DMR | -0.11 | $5.08 \times 10^{-03}$ | DMR |
| *KCNG2* | -0.08 | $4.74 \times 10^{-19}$ | DMR | -0.05 | $1.33 \times 10^{-08}$ | DMR |
| *RNF157* | -0.14 | $5.94 \times 10^{-12}$ | DMR | -0.42 | $1.34 \times 10^{-02}$ | DEG |
| *PPIAL4G* | -0.15 | $9.66 \times 10^{-11}$ | DMR | -0.24 | $5.36 \times 10^{-15}$ | DMR |
| *PTPRU* | 0.24 | $2.43 \times 10^{-06}$ | DMR | 0.11 | $7.68 \times 10^{-11}$ | DMR |
| *CHSY1* | 0.10 | $4.88 \times 10^{-05}$ | DMR | 0.08 | $8.27 \times 10^{-05}$ | DMR |
| *STX2* | -0.06 | $6.29 \times 10^{-05}$ | DMR | -0.13 | $2.70 \times 10^{-03}$ | DMR |
| *TRIM36* | -0.95 | $2.78 \times 10^{-04}$ | DEG | 0.12 | $1.03 \times 10^{-02}$ | DMR |
| *KLK4* | 0.16 | $8.52 \times 10^{-04}$ | DMR | 0.09 | $1.44 \times 10^{-03}$ | DMR |
| *NCOR2* | -0.08 | $1.11 \times 10^{-03}$ | DMR | -0.08 | $1.37 \times 10^{-03}$ | DMR |
| *SBK1* | -0.10 | $1.19 \times 10^{-03}$ | DMR | 0.10 | $1.54 \times 10^{-05}$ | DEG & DMR |
| *KIFAP3* | 0.03 | $1.35 \times 10^{-03}$ | DMR | 0.05 | $8.88 \times 10^{-03}$ | DMR |
| *GAS6* | -0.22 | $2.77 \times 10^{-03}$ | DMR | -0.06 | $8.21 \times 10^{-03}$ | DMR |
| *HAPLN4* | 0.15 | $3.58 \times 10^{-03}$ | DMR | 0.10 | $2.02 \times 10^{-04}$ | DMR |
| *RAB12* | -0.13 | $5.24 \times 10^{-03}$ | DMR | -0.08 | $1.90 \times 10^{-03}$ | DMR |
| *TAF1B* | 0.11 | $6.58 \times 10^{-03}$ | DMR | 0.12 | $1.84 \times 10^{-16}$ | DMR |
| *ADGRD1* | 0.10 | $7.40 \times 10^{-03}$ | DMR | 0.10 | $2.84 \times 10^{-03}$ | DMR |
| *SGTA* | 0.07 | $1.73 \times 10^{-02}$ | DMR | 0.09 | $3.24 \times 10^{-02}$ | DMR |
| *ZBTB7A* | -0.12 | $2.30 \times 10^{-02}$ | DMR | -0.83 | $3.62 \times 10^{-02}$ | DEG |
| *CEP170* | -0.58 | $3.98 \times 10^{-02}$ | DEG | -0.15 | $4.78 \times 10^{-24}$ | DMR |
| *AURKC* | 0.16 | $4.21 \times 10^{-02}$ | DMR | 0.10 | $6.35 \times 10^{-05}$ | DMR |
| *NUDT10* | 0.82 | $4.29 \times 10^{-02}$ | DEG | 1.00 | $3.82 \times 10^{-02}$ | DEG |
| *MEIS2* | -0.11 | $4.94 \times 10^{-02}$ | DMR | -0.09 | $1.76 \times 10^{-02}$ | DMR |
| *FBXO31* | - | - | - | 0.09 | $5.73 \times 10^{-03}$ | DEG &<br>DMR |
| *LTBR* | - | - | - | 0.12 | $1.88 \times 10^{-16}$ | DEG &<br>DMR |
| *ZNF423* | - | - | - | 0.07 | $3.90 \times 10^{-03}$ | DEG &<br>DMR |
| *AATF* | - | - | - | 0.12 | $3.02 \times 10^{-02}$ | DEG &<br>DMR |

To explore if Ov-HRT treatment modulates the effect of Ov, we also computed an interaction test between Ov-HRT treatment and Ov status. As expected for the sample size and the nature of combining CpGs to build DMRs, FDR adjustment left no DMRs. *LTBR*, as mentioned above, is a DE gene that is also a significant DMR in the PFC (Figure 3). The DMR contains 57 DMCs, and maps to the promoter and the first alternative exons of this gene. The DMR is significantly hypermethylated in Ov samples as compared to OI samples (average DNAm = 27% vs 14%; respectively), while in Ov-HRT the methylation level was not different to OI or Ov (average DNAm = 20%). The promoter/enhancer assays and the position of the DMR suggest that this DMR is functioning as a promoter (Figures S1). In agreement with this DMR functioning as a promoter, we observed a downregulation of *LTBR* with the hypermethylated DMR in Ov (Figure 3), while the expression in OI and Ov-HRT did not differ.

**FIGURE 3.**
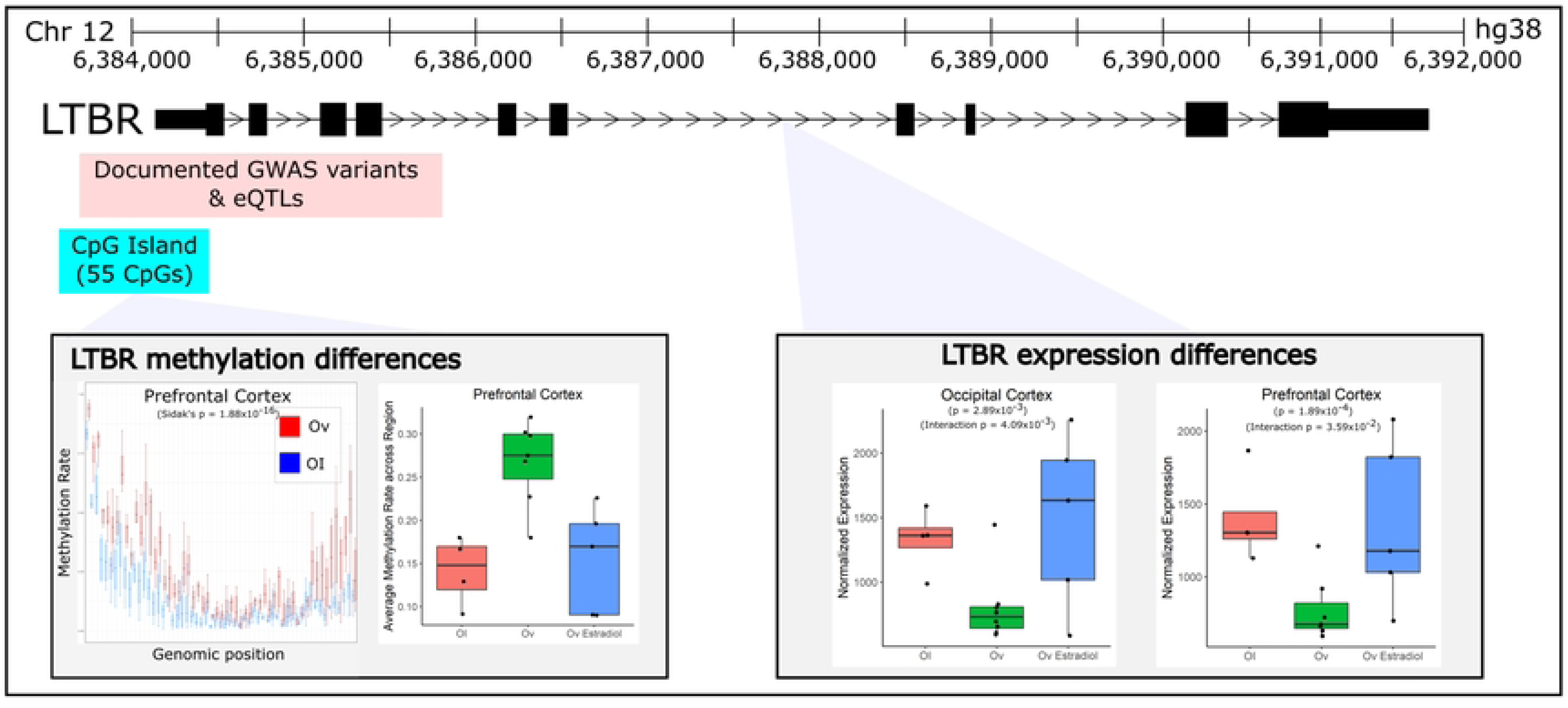
Ovariectomy-related changes in gene expression and methylation of *LTBR*. *LTBR* was a gene that demonstrated differences in expression associated with ovariectomy across both brain regions. In addition, it demonstrated significant differences in methylation in the PFC (but no significant DMR was identified in the OC). The DMR and CpG island that was annotated to *LTBR* resides just upstream of *LTBR* and appears to be hyper-methylated with ovariectomy. With average methylation rates, it appears that the hyper-methylation in this region that is caused by Ov is modulated by estradiol treatment. Corresponding expression of *LTBR* decreases with ovariectomy, but appears to be rescued to some degree with E2 treatment, particularly in the Occipital cortex. Near the same region of the DMR, lie a number of cis-acting eQTLs associated with *LTBR* expression that are also known GWAS variants for a variety of disorders – many of which are related to immune-disorders.

### 2.4 | Network analysis

Network analysis for each brain region was completed independently for each tissue. In the OC we identified 150 DE genes and 254 DMRs, while we identified 128 DE genes and 457 DMRs significantly associated with Ov in the PFC (Table S1). In the OC, 23 genes clustered into 3 MCODE clusters (MCODE score > 4.0) and in the PFC, 48 genes clustered into 5 MCODE clusters (MCODE score > 4.0) (Figures S2 & S3). Pathway analysis shows different pathways enriched in a tissue-specific manner, highlighting the particular function of each brain region. For instance, in the OC there was an enrichment in the regulation of insulin secretion, with *UBE2M* and *ADM* being hypermethylated and the estrogen receptor *ESR2* being hypomethylated in Ov. In the PFC, there was an enrichment in GPCR signaling, which included the pro-opiomelanocortin *POMC*, the G protein signaling 17 (*RGS17*) or the G protein subunit gamma 2 and 7 (*GNG2* and *GNG7*), among others. The ankyrin signaling pathway and the HSP90 chaperone mediated activation of steroid hormone receptors were also enriched in the PFC with Ov. These pathways included *ANK1* and *ANK3*, several dyneins (*DYNLL2*, *DYNLT3* and *DYNC1LI2*), and the FK506-binding protein 4 (*FKBP4*).

Given that the different brain regions under study play critical roles in processing cognitive functions, after filtering and manual curation to identify genes that were replicated across brain regions for omics, 891 total genes were submitted for a global integrated network analysis (Figure 1). Of the 891 genes, 127 genes clustered in 7 MCODE clusters (MCODE score > 4.0). The largest of the clusters contained 67 genes and was strongly enriched for “MAPK signaling”. Genes with differential expression and methylation from both brain regions are similarly represented in this pathway. For instance, and within the regulators of G protein signaling (RGS) subcluster, all three members of the R12 family (*RGS10*, *12* and *14*) were hypermethylated in Ov in the PFC. Exon 24 of the *RGS6* was downregulated in the OC leading to the production of different RGS6 transcripts under Ov conditions (Figure S4). This brain specificity in expression of RGS members may suggest activation/inhibition of different intracellular signaling cascades by brain region. One of the strongest results from the DNAm analyses, *UBE2M*, appears to play a key role in the regulation of this network (Figure 1). *UBE2M* interestingly links into the pathway through 17β-estradiol receptor, *ESR2*, which is hypermethylated in Ov in the OC. Other key networks worth highlighting include those that were enriched for “AMPK signaling” and several signaling pathways involved in transcription and translation regulation (i.e. ribosome biogenesis; Figure 1). Interestingly, an enrichment in “Dopaminergic synapse” was, almost exclusively, found in the PFC. Within this network, the family of kinesin motor proteins, KIF, were primarily downregulated (*KIF5A-C*; *KIF1B*, *KIF4A*, *KIFAP3*) in Ov. Among this family, *KIF26B* was hypermethylated in Ov, but DNAm levels went down with Ov-HRT, more similar to the levels in OI.

Using the 26 unique annotated genes from the interaction analysis across both brain regions, pathway enrichment suggests only two potential networks, HIF-1 signaling (*p*=0.0088, *LTBR*, *TIMP1*) and neuroactive ligand-receptor interaction (*p*=0.0096, *P2RX1*, *NMU*, *ADM*), suggesting that these genes and pathways may be working together in mediating the effects of HRT in Ov (Table 1).

28 genes showed overlap in consistent directions either across the two brain regions or both omics. These genes replicated across both regions or showed significant methylation and expression effects (Table 2). The network analysis highlights *UBE2M*, *AURKC*, *SGTA*, *RAB12*, *KIFAP3*, *NCOR2*, *TAF1B*, *ZBTB7A*, and *KCNG2* as genes with potentially key biological importance related to Ov across both brain regions (Figure 1).

## 3. | DISCUSSION

### 3.1 | Overall goals

This study examined the molecular effects of estrogen depletion at an older age on two different cortical regions, the OC and PFC, implicated in cognitive function. The OC controls visuospatial processing, distance and depth perception, color determination, object and face recognition, and memory formation [53, 54]. While the PFC is involved in working memory, temporal processing, decision making, flexibility, and goal-oriented behavior [52]. Damage to these brain regions contributes to cognitive decline in dementia patients. While limited in sample size, this study leveraged an NHP model that was sufficiently powered to detect significant differences in gene expression and DNAm because of our ability to tightly regulate the environment and obtain high quality and highly reproducible brain samples. With this NHP model of middle-aged female rhesus macaques, we identified highly translatable molecular changes in the brain that are linked with the E2 depletion associated with Ov. Given the natural depletion of E2 that occurs with age, these results present a novel understanding of the role E2 plays in the aging brain and how long-term immediate HRT treatment (∼4 years) can reverse or palliate those changes to maintain the brain in an age-matched molecular profile.

Because of the established relationship between E2 levels, its broad molecular regulatory function, and cognition [20, 30], we expected to identify robust molecular differences in these cognitive-relevant brain regions associated with Ov. For this reason, we completed RNA-sequencing to determine the genes that were DE and genome-wide DNAm sequencing to determine the genomic regions that were differentially methylated in the brains of animals that had undergone Ov for ∼4 years prior to necropsy. As expected, we detected a large number of DMRs as well as DE genes – sometimes genes that were both DE and differentially methylated in their respective promoter/enhancer regions (Figures 1-2). We note, however, that because these results are derived from heterogeneous bulk tissue that contains many cell types, we are unable to attribute these differences to actual changes in the cell-type molecular mechanisms linked to Ov. Instead, it is possible the changes we identify are driven by differences in the proportions of particular cell types. For example, *LTBR* is a gene that is primarily expressed in microglial cells. The differences we see in DNAm and expression between groups may either be attributed to a change in the abundance of microglia seen between groups or an actual shift in DNAmlevels across cell types (or even just a large DNAm shift in microglia). A follow up study using single-cell RNA-Seq would need to be completed to determine what the real drivers are in most of these cases.

### 3.2 | Ov is associated with dramatic changes in neural signaling pathways

Pathway analysis revealed several networks of genes that changed with Ov (Figure 1) in both brain regions. Importantly, all the identified networks had at least one gene that was DE and/or differentially methylated in both brain regions, suggesting a common link to the same pathways across them. These include the voltage-gated potassium channel encoded by *KCNG2* (proteasome, p=2.53×10^-2^), the TATA-box binding protein associated factor (*TAF1B*, ribosome biogenesis, *p*=8.41×10^-6^), the nuclear receptor corepressor 2 (*NCOR2*, NOTCH signaling, *p*=2.65×10^-2^), the Ras-related protein 12 (*RAB12*, AMPK signaling, *p*=2.96×10^-2^), and the kinesin associated protein 3 (*KIFAP3*, dopaminergic synapse, *p*=2.37×10^-6^). Interestingly, *KIFAP3* was downregulated in the PFC of Ov-HRT females under a chronic obesogenic diet [55]. In the current study, *KIFAP3* was hypermethylated in the PFC with Ov as compared to OI, while the DNAm levels were similar to the group receiving HRT. These results highlight the role of this gene in the PFC and its responsiveness to the presence of E2, independently of the diet. Within the basal transcription factors network (*p*=4.43×10^-4^), the zinc finger and BTB domain containing 7A factor (*ZBTB7A*) is known to transcriptionally upregulate ERα expression by directly binding to the *ESR1* promoter. In addition, ERα potentiates *ZBTB7A* expression via a positive loop in breast cancer [56]. In the PFC, *ZBTB7A* was downregulated, probably due to the absence of E2 in Ov, and, given the role of this transcription factor in metabolism [57], this downregulation could have implications in the regulation of brain metabolism.

The dopaminergic synapse pathway contained the *KIFAP3* gene which was hypermethylated in both brain regions and contained a number of other members of the kinesin heavy chain proteins (i.e. *KIF1B*, *4A*, *5A, 5B* and *5C* were all downregulated in the PFC). Kinesins are molecular motors that transport cargos along microtubules [58]. These kinesin members are involved in transporting mitochondria, amyloid precursor protein vesicles, GABA and dopamine receptors, lysosomes, choline acetyl transferase and dopamine [59–65]. Our results suggest that with Ov there is a downregulation in kinesin expression that could be contributing to alterations in intracellular protein trafficking that could impact synaptic transmission.

One of the strongest biological pathways enriched in Ov was the MAPK signaling (enrichment *p*=2.3×10^-10^). Prior evidence showed that E2 alters cellular components required for maintaining balance between active and inactive MAPKs. For example, crosstalk between phosphorylation and ubiquitination pathways can exert long-term changes in cellular processes through multiple feedback loops that ultimately impact apoptosis and cell proliferation [66]. Among the members of the MAPK signaling pathway, the ERK 1 gene (*MAPK3*) and the ubiquitination gene *UBE2M* were hypomethylated, and several RGS proteins (*RGS 10, 12* and *14*) were hypermethylated in the PFC with Ov. Activation of ERK 1/2 subjected to G-protein coupled receptor-mediated signaling is regulated through RGS proteins [67, 68]. Although additional studies analyzing protein levels and activation/inhibition ratio of these molecules are needed, our results suggest that RGS protein activity might be downregulated, leading to less inhibition of ERK 1, which would be consequently upregulated (supported by the hypomethylated DMR mapping to *MAPK3*). In addition, three genes are differentially methylated in both brain regions, *UBE2M*, *SGTA* and *AURKC*. While little is known about the neural function of aurora kinase C (*AURKC*), the *SGTA* (small glutamine-rich tetratricopeptide repeat-containing protein alpha) encodes for a molecular co-chaperone that interacts with steroid receptors and heat shock chaperone proteins, i.e. HSP90AA1 (Figure 1), to regulate steroid receptor signaling, protein folding and conformation state, receptor stability, subcellular localization and intracellular trafficking [69]. A study in yeast showed that Hsp90 functions to maintain the estrogen receptors in a high affinity hormone-binding conformation [70]. Interestingly, *UBE2M* involvement in the MAPK signaling network is through its interaction with the ERβ gene (*ESR2*), that was hypermethylated in Ov (Figure 1). Given its robust association with Ov across both brain regions and both the transcriptome and methylome, *UBE2M* is a strong result that should be heavily considered for further investigation. We confirmed that the DMR proximal to *UBE2M* functions as a promoter (Figure S1), suggesting that changes in DNAm in this DMR may contribute to regulating its expression. UBE2M’s primary function is as a ubiquitin-protein transferase, involved in protein neddylation, which is a post-translational ubiquitin-like protein modification that plays pivotal roles in protein quality control and homeostasis. Neddylation, involving UBE2M and other enzymes, is a critical mechanism for targeting and degrading misfolded or damaged proteins, helping to maintain protein quality control within brain cells [71]. For instance, during the initial stages of AD, ubiquitin-proteasome proteolysis degrades the abnormal amyloid β peptides and hyperphosphorylated tau. But as the disease progresses, ubiquitination becomes ineffective at degrading the accumulating insoluble proteins, and neddylation seems to contribute to degradation with these abnormal proteins. In AD patients, neddylation mechanisms are dysregulated [72], and neurons show accumulation of the neddylation enzyme NEDD8 in the cytoplasm and colocalization with ubiquitin and proteasome components in protein inclusions in the brain [72, 73]. Our results showed hypomethylation in both brain regions and upregulation of *UBE2M* in the OC. While additional studies are needed to determine the cellular localization of *UBE2M* and its role in protein neddylation, our results suggest that dysregulation of *UBE2M* associated with E2 depletion may be a key player mediating the negative effects that lack of E2 has on brain function [74]. Together, these results emphasize *SGTA* and *UBE2M,* through their direct (*ESR2)* or indirect connections with the estrogen system, as critical mediators of the MAPK signaling cascade and its connections with synaptic function, neuroinflammation and neurodegeneration across brain regions [75–78].

### 3.3 | Immediate estradiol supplementation ameliorates molecular alterations linked to Ov

After identifying molecular changes associated with Ov, we were interested in understanding whether immediate (right after Ov) and long-term (over 4 years) E2 treatment reverses any of the changes linked to Ov. By identifying Ov-linked effects ameliorated by E2 treatment, the biological pathways that are directly impacted by estrogen levels become clearer. While E2 treatment ameliorates some of the behavioral and physiological changes seen following menopause in humans, the effects of E2 treatment on cognitive performance are still mixed [18]. Such inconclusive results could be due to the differences in HRT timing, diet, and other confounding variables common in human studies. Contrarily, results in NHPs on the beneficial effects of HRT on cognition are more consistent [30, 41], probably due to the controlled experimental conditions. Thus, by understanding the specific genes altered by E2 in the brain with this animal model, we can begin to understand the biological pathways estrogen impacts and thereby develop more targeted therapeutics that specifically improve brain function and ultimately cognitive performance.

In the OC and PFC, we identified 19 and 10 genes, respectively, with suggestive evidence for E2 effects, where E2 appears to restore expression levels to a level that is similar to OI (Table 2). Among these 29 genes, the following are known to interact with the estrogen receptors, or its expression being associated with the levels of estrogen. The transient receptor potential vanilloid 6 (*TRPV6*) is a highly Ca^2+^-selective channel that contains an ERE in its promoter [79], and its regulation by estrogen has been proposed in peripheral tissues [80] and the CNS, including cortex [81]. Furthermore, in mice, hypothalamic levels of Trpv6 are susceptible to estradiol oscillations through the estrous cycle, with higher Trpv6 levels at the proestrous phase where estrogen levels are at their highest [81]. In the OC, *TRPV6* was upregulated with Ov, and the levels decreased with HRT. These results disagree with previously reported findings in the hippocampus [81] and could stem from brain-specific differences in *TRPV6* regulation. Adrenomedullin (*ADM*) is a peptide exerting important functions in the periphery and CNS. In the uterus, studies revealed that ADM promoter is recognized by the ER in a ligand-dependent manner, and that there is a positive correlation between estrogen levels and *ADM* gene expression [82–85]. In the CNS, ADM is known to contribute to the activation of the hypothalamic-pituitary-adrenal (HPA) axis through release of CRH [86], and thus contributing to regulating hormonal responses to stress. In breast cancer cell lines, the insulin-sensitive glucose transporter 12 (*SLC2A12*) protein levels are increased with estradiol [87]. The fucosyltransferase 1 (*FUT1*) is involved in the creation of a precursor of the H antigen, which is required for the final step in the synthesis of soluble A and B antigens, and its expression is mediated by the ESR2 [88]. Neuromedin U (*NMU*) is a gonadal peptide, which receptor, neuromedin U2 is expressed throughout the brain [89]. It is suggested that *NMU* has a protective role in neurodegenerative diseases [90]. In particular, *NMU* protects neuronal cell viability, and inhibits inflammation-induced memory impairments [90]. Others have shown that estradiol levels following Ov in rats resulted alterations in *NMU* levels in the brain [91, 92]. Although this effect seems to be estradiol dose-dependent, with low estradiol levels increasing *NMU* expression levels, while this increment was only previously observed with progesterone supplementation [91, 92]. The Purkinje cell protein 4 (*PCP4*) is a calmodulin-binding anti-apoptotic peptide in neural cells and an estrogen-inducible peptide in breast cancer cell lines [93]. The hematopoietic PBX-interacting protein 1 (*PBXIP1*) is a transcription factor involved in extracellular matrix organization and chromatin regulation. It is an estrogen receptor (ER) interacting protein that regulates estrogen-mediated breast cancer cell proliferation and tumorigenesis [94]. *PBXIP1* is a key gene in the AMPK signaling network that was altered by Ov (Figure 1). While the specific function of these genes in the cortex, or even in the brain, remains to be investigated, our results indicate that Ov and E2 treatment alter these genes (and likely associated pathways) to normalize them to those levels of age matched controls.

Enrichment analysis of these 29 DE genes revealed two small networks that were significant: HIF-1 signaling (*p*=0.0088, *LTBR*, *TIMP1*) and neuroactive ligand-receptor interaction (*p*=0.0096, *P2RX1*, *NMU*, *ADM*). HIF-1 signaling has been previously linked to estradiol [95] and has been shown to regulate neuroinflammation in traumatic brain injury [96]. This network contains the lymphotoxin B receptor (*LTBR*), which is one of our most promising results that demonstrates differential expression effects in both brain areas. We also identified a DMR mapping to *LTBR* in the PFC (Figure 2). We confirmed that the DMR proximal to *LTBR* is a promoter (Figure S2). LTBR has been mostly studied in lymphoid tissues, with data supporting its role as a regulator of inflammation [97], a process known to be modulated by estrogen as well [98]. In fact, LTBR signaling can activate both canonical and alternative NF-κβ signaling to induce proinflammatory chemokines and cytokines [99]. Although studies on the role of *LTBR* in brain function remain mostly unexplored, a recent transcriptomic study identified *LTBR* as a neuroinflammatory biomarker of AD [100]. More specifically, it has been recently shown that *LTBR* is upregulated in microglia of aged (mean age of 94 years) brains as compared to middle-aged (mean age of 53 years) human brains [101], which could suggest that upregulated *LTBR* levels might be mediating age-related neuroinflammation through microglial function. However, the relationship between *LTBR* expression/function and microglial activation status remains unknown, and additional studies are needed to determine how or whether *LTBR* mediates microglial activation. Others have shown that estrogen can inactivate microglia through the ERβ [102]. Our results show increased DNAm levels and decreased levels of *LTBR* expression with Ov (Figure 2), with *LTBR* expression levels reverting to OI levels with HRT. As discussed earlier, it is possible that the changes in DNAm and gene expression seen with Ov-HRT are associated with a change in the proportions of cell types, namely microglia, with Ov-HRT. This is a limitation in this study, and we cannot exclude that such molecular changes are due to a change in the number of microglia present, their activation status or a combination of all. Nonetheless, it is evident from our results that understanding the role of *LTBR* in microglia function would be of critical importance to understanding estrogen’s relationship to brain function.

## 4 | CONCLUSIONS

This work highlights the importance of multiple omics across multiple brain regions to identify robust molecular signals linked to E2 regulation in the aging brain. By including multiple omics and more than one brain area, we were able to home in on biological effects that replicated across brain regions and thereby reduce noise from false positives. Importantly, this work represents a major step towards understanding molecular changes in the brain that are linked to Ov and how HRT may revert or protect against the negative consequences of a depletion in E2. Our findings indicate that the molecular profile of the cortical regions (OC and PFC) in the absence of E2 may lead to neuroinflammation or the dysregulation of critical processes, like intracellular axonal protein trafficking or protein ubiquitination, that are necessary for proper function of brain cells. Immediate HRT reverted these effects, at least partially, by bringing the epigenetic/transcriptomic profile of genes involved in neuroinflammation and other unknown functions, to that profile of age-matched OI females. It remains to be known if the molecular profile of the brain after HRT is more similar to that of younger OI brains, which we are currently investigating. Nonetheless, our results present real opportunities to discover novel therapeutics to slow cognitive decline caused by the lack of E2. Although our work needs further validation with larger cohorts, it also requires focused investigation of some of the genes we identified with very robust effects like *LTBR* and *UBE2M*. Moreover, because our studies were performed in a NHP preclinical animal model of human aging, the findings may have more immediate translational potential to clinical studies involving postmenopausal women.

## 5 | MATERIALS AND METHODS

### 5.1 | Subjects

This study was approved by the Oregon National Primate Research Center (ONPRC) Institutional Animal Care and Use Committee and used 19 old (range = 15.4-19.2 years, at the beginning of the study) female rhesus macaques (*Macaca mulatta*). The maximum lifespan of this species is in the early 40s, so these animals were proportionally in late-middle to early-old age [103] and in the range of pre- to peri-menopausal endocrine status [40, 104]. The animals were socially housed indoors in paired cages under controlled environmental conditions: 24 °C temperature; 12-h light and 12-h darkness photoperiods (lights on at 07:00 h) and were cared for by the ONPRC Division of Comparative Medicine in accordance with the National Research Council’s Guide for the Care and Use of Laboratory Animals. Daily meals at ∼08:00 h and ∼ 15:00 h were supplemented with fresh fruits or vegetables; fresh drinking water was available ad libitum. Diet was monkey chow which provides calories with 13% fat, 69% complex carbohydrates (includes 6% sugars), and 18% protein. Additional enrichment included watching video programs and interactions with the Behavioral Science Unit staff and animal care technicians.

### 5.2 | Ovariectomy and estradiol supplementation

Before ovariectomy, all of the females were showing menstrual cycles and were therefore considered to be premenopausal at the beginning of the study. Except for the ovary intact (OI, *n* = 4) females, the rest of the animals were Ov, resulting in E2 levels below 20 pg/mL. Half of the females (*n* = 6, Ov-HRT) were immediately started on HRT in the form of E2-containing elastomer capsules, which achieved serum E2 concentrations of 94.3 ± 20.5 pg/ mL; the other half (*n* = 8) received empty capsules (placebo), which achieved serum E2 concentrations of <30 pg/mL on average across ∼48 months (age at end of study, 19.4–23.2 years). Serum E2 was measured every 2 months and the capsule replaced or its size adjusted as deemed appropriate [30].

### 5.3 | Euthanasia

After the ∼4-years duration of the study, a detailed necropsy protocol previously used in our laboratory was used to systematically collect brain tissues from all subjects; other body tissues were made available to other investigators for unrelated postmortem studies. Briefly, monkeys were sedated with ketamine (10 mg/kg), and administered pentobarbital, followed by exsanguination, as recommended by the 2013 Edition of the American Veterinary Medical Association Guidelines for the Euthanasia of Animals. Brains were quickly removed and the right hemisphere was dissected to isolate the different brain regions. Briefly, the dorsal and ventral banks of the dorsolateral prefrontal cortex (PFC) were collected around the primary sulcus. The OC was removed from the caudal tip of the occipital lobe. All tissues were wrapped in aluminum foil and immediately frozen in liquid nitrogen, and then archived at −80 °C.

### 5.4 | DNA/RNA isolation

Genomic DNA and RNA were extracted from each brain region using the All-Prep DNA/RNA/miRNA Universal kit (Qiagen Sciences Inc., Germantown, MD) following the manufacturer’s recommendations. Briefly, each brain region was pulverized and ∼30 mg of tissue was used for DNA/RNA isolation.

### 5.5 | cDNA library construction and sequencing

For stranded RNA-seq, cDNA libraries were prepared with the TruSeq stranded mRNA library prep Kit (cat# RS-122-2101, Illumina, San Diego, CA, USA). The resulting libraries were sequenced on a HiSeq 4000 (Genomics & Cell Characterization Core Facility, University of Oregon) using a paired-end run (2 × 150 bases). A minimum of 100 M reads was generated from each library.

### 5.6 | RNA-Seq Processing and Calling

Raw sequences were examined for quality using FASTQC [105]. Phred scores (probability a base was called correctly) and GC content were observed for abnormalities. After initial quality control of the reads was completed, alignment was performed using STAR two pass alignment [106]. Reads were aligned to the *Macaca mulatta* assembly (Mmul_10) and scored on how well they corresponded to the reference genome and whether or not they map to multiple positions across the genome. Low scoring reads, usually short and poor-quality reads, were not retained (Mapping Quality Score < 2). Post-alignment, reads were quantified at the gene level using the program featureCounts [107], and DESeq2 [108] was used to transform gene counts and estimate fold-change values for differentially expressed genes. Genes that had either an average read count below 5, missing values for more than one-third of the samples, or a coefficient of variation greater than 50 in the control samples were dropped to remove noisy and lowly expressed genes.

### 5.7 | Differential Expression Analysis

Differential expression analysis was computed in DESeq2 where the gene expression values were evaluated as the outcome in a negative binomial generalized linear model [108]. Whether or not the animal had undergone Ov was the predictor of the primary model we computed. This analysis was computed only in animals that had not received Ov-HRT treatment. Given no significant differences in age between animals and all of them are all much older females (∼25 yrs) in the same stage of life and the limited power of the study we did not adjust for age as a covariate. A Benjamini-Hochberg false discovery rate (FDR) was applied to the unadjusted p-value to account for multiple comparisons [109]. We repeated this same analysis in animals treated with OV-HRT. In addition to these models to find differences directly related to Ov, we tested the interaction between Ov-HRT and Ov. To enable this analysis, we replicated the control samples (OI) into two groups, one labeled as having received Ov-HRT treatment and the other as having not received Ov-HRT treatment. We recognize that comparing both Ov groups to the same exact set of controls (OI) will lead to false positives, but we primarily used the test of interaction as a way of rapidly identifying genes associated with Ov that are potentially modified by OV-HRT treatment. As expected, FDR adjustment left no interaction results, so we considered interaction results that met an unadjusted *p* < 0.05, particularly because computing the interaction globally for all genes was not our main interest. We were primarily interested in genes that showed expression changes related to Ov that no longer showed association with Ov with Ov-HRT treatment. To obtain a general set of genes that fit this criterion, we filtered the results to include only genes that showed suggestive evidence of Ov association (unadjusted *p* < 0.05) without Ov-HRT treatment and little evidence of Ov association with Ov-HRT treatment (unadjusted *p* > 0.1). From this reduced set of genes, we further filtered down to only those results that had at least suggestive evidence (*p* < 0.05) of interaction between Ov and Ov-HRT treatment.

### 5.8 | Differential Exon Usage Analysis

The DEXSeq pipeline was applied (with the default parameters) to analyze the aligned reads and obtain exon level counts [110]. The exon level counts were loaded and inspected in R (4.1.1) as DEXSeq objects before being normalized with DESeq2’s normalization algorithm [108]. DEXSeq, similar to DESeq2, computes negative binomial regression and shares dispersion estimation across features. The program is designed to estimate differences in exon usage within a particular gene across conditions and will not identify genes with global differences in exon expression across a given gene (i.e., the genes identified by DEU analysis will be different than those identified in the DE analysis). We computed a model where the predictors included dummy variable for the exon, an indicator variable for whether the animal had undergone Ov, as well as the interaction between the two to assess DEU age associated with Ov. Results with an FDR adjusted *p* < 0.05 were retained for pathway analysis. Top results were also overlapped with DMR genes to assess whether changes in methylation appeared to be affecting exon usage of any genes.

### 5.9 **| Genome-wide DNA methylation profiling**

Genomic DNA was checked for quality by electrophoresis on a 0.7% agarose gel, using a NanoDrop 8000 spectrophotometer (Thermo Scientific, Wilmington, DE, USA) and quantified using a Qubit (Thermo Scientific, Wilmington, DE, USA). Five hundred nanograms of genomic DNA was sheared using a Bioruptor UCD200 (Diagenode, Denville, NJ, USA), generating fragments ∼180 bp. The Illumina TruSeq Methyl Capture EPIC library prep kit (Illumina, Santa Clara, CA, USA) was used following the manufacturer’s instructions. The EPIC probes interrogate >3.3 million individual CpG sites per sample at single-nucleotide resolution. After end repair, 3’ A-tailing, and adaptor ligation, libraries were pooled in groups of four, followed by two rounds of hybridization and capture using the EPIC probes, bisulfite conversion and final amplification. After library quantification using a 2100 Bioanalyzer (Agilent Technologies), DNA libraries were sequenced (3 libraries per 150PE lane) on an Illumina HiSeq4000 at the University of Oregon Genomics & Cell Characterization Core Facility (GC3F). Five percent PhiX DNA (Illumina Inc.) was added to each library pool during cluster amplification to boost diversity. Cases and control samples were mixed within lanes and sequenced together on the same flow-cell to reduce the impact of batch effects on data. The quality of the bisulfite-converted sequencing reads was assessed with FastQC [105]. Reads were trimmed and aligned to the macaque reference genome (Mmul10), and then the bisulfite conversion rates were evaluated, insuring all libraries were >98% converted, and CpG methylation was evaluated using Bismark [111]. The methylation rates were calculated as the ratio of methylated reads over the total number of reads. Methylation rates for CpGs with fewer than 10 reads were excluded from further analysis. We next removed CpG sites on sex chromosomes. The remaining ∼2.8-3.0 million CpGs per sample (OC and PFC; respectively) post-filtering were used for downstream analyses.

All sequence reads were submitted to the Sequence Read Archive at NCBI under project accession number TBD.

The differential methylation analysis was carried out by applying a generalized linear mixed effects model (GLMM) implemented in R package PQLseq (version 1.2.1) [112, 113] separately for each CpG site. PQLseq models the technical sampling variation in bisulfite sequencing data with a binomial distribution, effects of biological and technical covariates with the linear model, and the random effects with a correlated multivariate normal distribution. The most common methylation proportion values are 0 and 1, which are problematic in the context of generalized linear models with the logit link function (infinite in the logit-transformed space). We used a common pseudo-count transformation to avoid both extremes, as recommended, for example, by the developers of PQLseq [114]. This was done after adding + 1 to the numbers of methylated reads and +2 to the total numbers of reads to avoid modeling methylation proportions that are exactly 0 or 1, as recommended by the authors of PQLseq [114]. This pseudo-count transformation was only applied to non-missing values (coverage >10×).

Each nominal p-value was corrected for multiple comparisons by the False Discovery Rate (FDR). In parallel, the nominal p-value was used as input for Comb-p [115] analysis to identify differentially methylated regions (DMRs) between controls and AUD subjects as previously described [116].

### 5.10 | Network analysis

Significant DMRs that had gene annotations, as well as DE genes and genes demonstrating DEU were combined for each brain region. Those gene lists were again combined across brain regions to identify significantly altered genes that replicated across both, OC and PFC. Genes that were DE in both tissues were only retained if they were altered in consistent directions across both brain regions.

Results were analyzed in KEGG, STRING, and MCODE to find biological pathways enriched between groups (Figure 1, Figure S2-S3, Table 1, Table 2) [117, 118]. STRING was used to obtain protein-protein interactions for all genes that met our filtering criteria for each omic analysis. STRING was applied to find only “High confidence” protein-protein interactions with options for “textmining” and “neighborhood” disabled [117]. MCODE was applied to the remaining interactions to obtain a set of highly interconnected gene clusters [118] and the biological functions of each clusters with MCODE scores greater than 4.0 were identified through the KEGG pathways [119].

### 5.11 | Functional promoter/enhancer assay

To determine the promoter or enhancer activity capacity of two DMRs located in the promoter and overlapping with exon 1 of the rhesus macaque *LTBR* and overlapping with the last exon of *MZF1* and in the promoter of *UBE2M* genes, we cloned the corresponding macaque DMR regions (*LTBR* (PFC), chr11:6528520-6529383; *UBE2M* (PFC), chr19:58128610-58130108 & (OC), chr19: 58128532-58130175) in the luciferase reporter vector pGL3 (Promega) and transfected HEK293 cells (HEK 293, obtained from the Wake Core Repository). In addition, we transfected cells with the basic and control pGL3 as negative and positive controls; respectively. HEK293 cells were seeded in 96-well plates at 10.000 cells/well density and cultured in Dulbecco’s modified Eagle medium (DMEM) containing high glucose (4.5 g/L) supplemented with 10% fetal bovine serum (FBS) and maintained at 37°C and 5% CO_2_. Twenty-four hours later, cells were transfected using 90ng of each corresponding vector diluted in 10ul of opti-medium and 0.3ul of X-treme GENE HP DNA Transfection Reagent (Roche). 10ng of Renilla vector (Promega) was co-transfected and used for normalization. After 48 hours of transfection, Dual-Glo® Reagent equal to the volume of culture medium was added to each well. After 10 minutes, firefly luminescence was measured in a luminometer (SpectraMax iD3).

## Grant support

National Institutes of Health Grants: P30 AG066518, P51 OD011092, RF1 AG062220

## Data Sharing

The data that support the findings of this study are available on GEO under the following accession number: TBD

## Author contributions

SGK, HFU and RCJ designed the experiments. SGK and HFU provided the rhesus macaque samples. RCJ isolated all the DNA and RNA samples, and prepared all the omics libraries. DZ and JB conducted the reporter assays. DNA methylation and gene expression bioinformatic analyses were performed by KDZ and LJW, and KDZ advised and conducted on the appropriate statistical analyses to be performed in all the experiments. SGK, HFU, KDZ and RCJ and RCJ wrote the manuscript with the help of all the authors.

## Conflict of Interest statement

The authors declare no conflict of interest

## Notes

### Competing Interest Statement

The authors have declared no competing interest.

